# Non-coding mutations reveal cancer driver cistromes in luminal breast cancer

**DOI:** 10.1101/2021.05.29.446210

**Authors:** Samah El Ghamrasni, Rene Quevedo, James Hawley, Parisa Mazrooei, Youstina Hanna, Iulia Cirlan, Helen Zhu, Jeff Bruce, Leslie E. Oldfield, S. Y. Cindy Yang, Paul Guilhamon, Jüri Reimand, Dave Cescon, Susan J. Done, Mathieu Lupien, Trevor J Pugh

## Abstract

Whole genome sequencing of primary breast tumors enabled the identification of cancer driver genes ^1,2^ and non-coding cancer driver plexuses from somatic mutations ^3–6^. However, differentiating driver and passenger events among non-coding genetic variants remains a challenge to understand the etiology of cancer and inform delivery of personalized cancer medicine. Herein, we reveal an enrichment of non-coding mutations in cis-regulatory elements that cover a subset of transcription factors linked to tumor progression in luminal breast cancers. Using a cohort of 26 primary luminal ER+PR+ breast tumors, we compiled a catalogue of ∼100,000 unique cis-regulatory elements from ATAC-seq data. Integrating this catalogue with somatic mutations from 350 publicly available breast tumor whole genomes, we identified four recurrently mutated individual cis-regulatory elements. By then partitioning the non-coding genome into cistromes, defined as the sum of binding sites for a transcription factor, we uncovered cancer driver cistromes for ten transcription factors in luminal breast cancer, namely CTCF, ELF1, ESR1, FOSL2, FOXA1, FOXM1 GATA3, JUND, TFAP2A, and TFAP2C in luminal breast cancer. Nine of these ten transcription factors were shown to be essential for growth in breast cancer, with four exclusive to the luminal subtype. Collectively, we present a strategy to find cancer driver cistromes relying on quantifying the enrichment of non-coding mutations over cis-regulatory elements concatenated into a functional unit drawn from an accessible chromatin catalogue derived from primary cancer tissues.

## Introduction

Breast cancer is the second leading cause of death in women in North America ^7^. Currently, treatment decisions rely on the histology and the expression of three proteins: estrogen receptor (ER), progesterone receptors (PR), and HER/neu (ERBB2) ^7^. Approximately 80% of all breast cancers are of the luminal (ER+) subtype, 65% of which are also PR+; together, the ER+PR+ luminal subtype makes up 52% of all breast cancers ^8,9^. Large-scale analysis of whole genome sequencing in breast tumors has identified 99 driver genes with recurrent protein-coding alterations ^1,2^ as well as a high number of mutations within the non-coding genome ^1^. Non-coding mutations can alter the transcription factor binding to the DNA and affect enhancer-promoter interactions to perturb gene expression ^3,10–19^. However, the inclusion of non-coding mutations to find cancer drivers remains a challenge in ER+PR+ luminal breast cancer that needs to be addressed to comprehensively resolve the role of genetic variants in oncogenesis.

The non-coding genome is known to harbor many of cis-regulatory elements, defined as binding sites for transcription factors involved in transcriptional regulation by serving as promoters, enhancers or anchors of chromatin interactions ^20^. In luminal breast cancer, cis-regulatory elements are bound by key transcription factors, including ESR1, FOXA1 and GATA3 which have a role in maintaining the luminal phenotype as well as the growth and differentiation of breast epithelium^21^. Disruption of either of these transcription factors or their binding sites can affect their binding to the chromatin^22^, which can modulate downstream gene expression. A subset of transcription factors active in luminal breast cancer are known as driver genes due to positive selection of protein-coding mutations ^23–25^.

Mutations within regulatory elements of enhancers and promoters can be responsible for the development of disorders with the same magnitude as mutations affecting protein-coding genes ^10–14,26,27^. A classic example of this is the *TERT* promoter which is frequently mutated across several cancer types as a mechanism for telomerase reactivation ^28^; it has been observed in 71% of sporadic melanoma and 60-75% of glioblastomas ^10–14,26,27^. Variants within the *TERT* promoters also lead to an increased risk of breast and ovarian cancer development ^29^. Pan-cancer analysis of the PCAWG project showed that the long tail of infrequent non-coding mutations in promoters and distal regulatory elements converged to pathways and molecular interaction networks of oncogenic processes ^19^. Zhu et al found frequently mutated regulatory elements in cancer genomes that interact with target genes via long-range chromatin interactions^19^.

The sum of all regulatory elements bound by a transcription factor in a given cell-type has been referred to as a “cistrome”^30^. Analysis of mutations across the cistromes of prostate cancer revealed a high frequency of mutations within the binding sites of key transcription factors including FOXA1, HOXB13 and AR ^17^. In luminal breast cancer, Bailey *et al*. found 7 functionally validated mutations within the cis-regulatory elements of *ESR1* that altered gene expression ^3^, while Cowper-Sal·lari *et al*. found that risk-associated SNPs in the cistrome of FOXA1 modulated the expression of downstream target genes ^22^. These studies highlight the key, albeit underappreciated role that cis-regulatory elements and cistromes play in tumorigenesis.

Within this study, we drew parallels to the approaches to finding driver mutations between the coding and non-coding genome, by defining cancer drivers as units of the genome that are enriched in mutations more than expected by chance. Similar to looking for hotspots of mutations within individual exons, we first focused on individual cis-regulatory elements across accessible chromatin regions. Proceeding to a broader scale, akin to looking at multiple exons that make up a gene, we explore for mutations across the cistromes of transcription factors in accessible chromatin regions of luminal breast cancer. Together, our study identified mutations clustered within cistromes of transcription factors essential to luminal breast cancer.

## Results

### Comprehensive chromatin accessibility analysis in primary ER+PR+ luminal breast cancer

To identify cis-regulatory elements, we used ATAC-seq to map the accessible chromatin of 26 luminal primary ER+PR+ invasive ductal carcinomas breast tumors freshly collected at the Princess Margaret Cancer Centre (PM_Lum; n=26) (**Table S1**) ^31,32^. To enrich for malignant cells, we used flow cytometry to sort cells from dissociated tumors using the anti-CD45RO (anti-CD45) antibody (**Figure 1a**). In the immune-depleted (CD45-) cancer cells, we identified a catalogue of 99,516 (41.37Mb) unique cis-regulatory elements found in accessible chromatin as defined by ATAC-seq peak coverage called using MACS2 ^33^ (**Table S2**).

**Figure 1:**
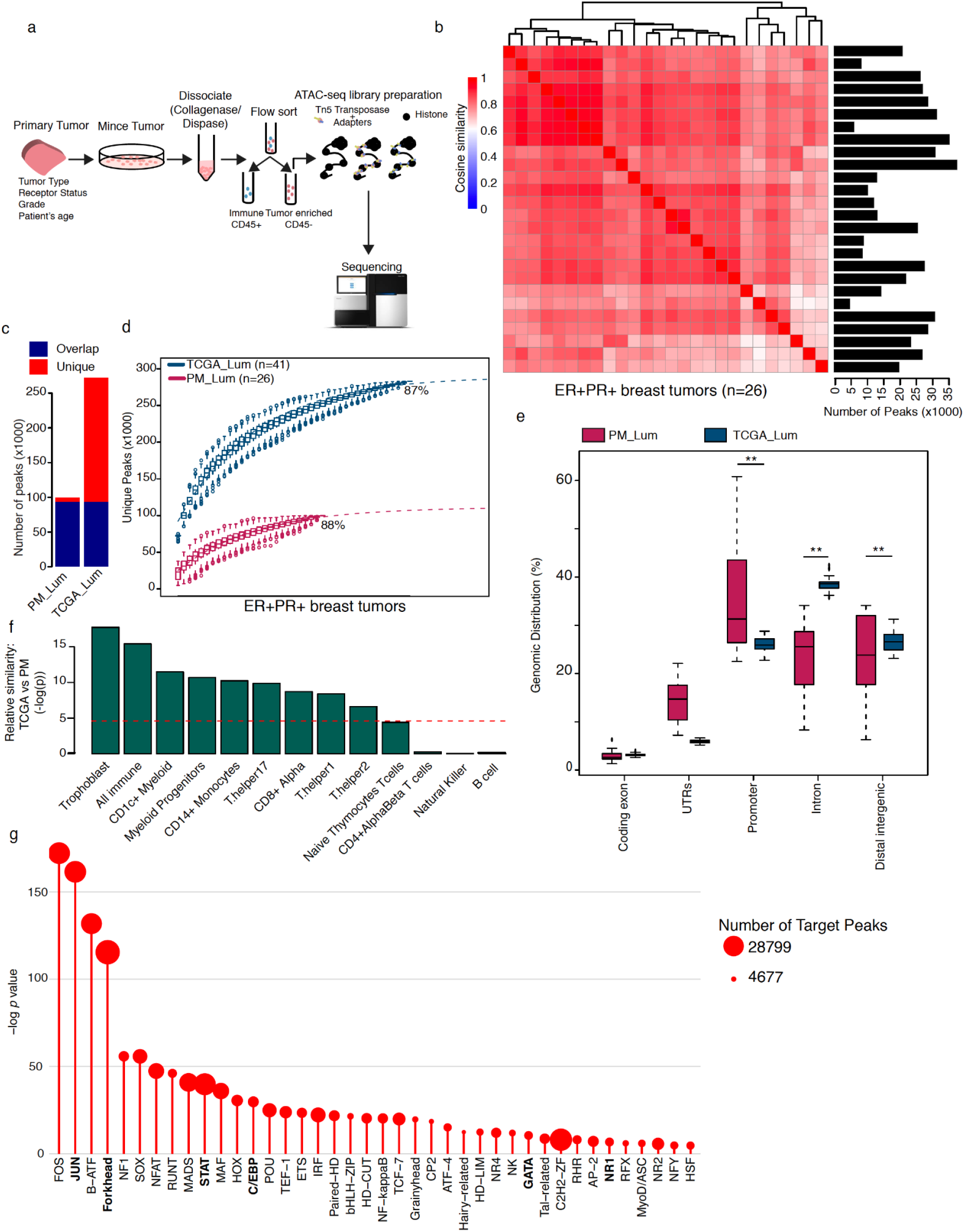
Identifying chromatin accessibility in ER+PR+ breast cancer. (**a**) Primary tumors were minced and dissociated for subsequent flow sorting into immune and epithelial cell populations, followed by ATAC-seq profiling. (**b**) Heatmap showing similarities between ER+PR+ open chromatin profiles. Cosine similarity analysis was calculated using comparing all chromatin accessibility of samples to each other. Barplot showing number of called peaks per sample. (**c**) Barplot showing the number of accessible chromatin regions from TCGA_Lum datasets that overlapped PM_Lum in blue and the ones unique to each cohort in red (**d**) A graph showing the chromatin accessibility saturation curve. A non-linear regression model analysis was performed using the number of unique ATAC peaks discovered in each sample to estimate the percentage of open chromatin mapped in PM_Lum (Purple; n=26 samples) and TCGA_Lum (Blue; n=41 samples). (**e**) Percentage of distribution of mapped open chromatin regions within the genome. The cis-regulatory element annotation system (CEAS) is utilized to perform genomic distribution analysis of the open chromatin region mapped by ATAC-seq. **pvalue < 0.001 (**f**) barplot showing p-values for cosine similarities between PM_Lum and TCGA_Lum in comparison to immune cells’ accessible chromatin. Red dotted line represents t-test p-value=0.01. (**g**) Lollipop graph showing enriched motif families in ER+PR+ breast tumors (p-value < 0.01). The catalogue of 26 ATAC-seq data was used. Enrichment of motifs within ATAC-seq regions against DNaseI hypersensitive sites from several cell lines was computed. Motif families were obtained using the Jaspar database. The size of the circles represents the number of target peaks for each motif.

To examine the quality of our data, we ran a similarity pairwise-comparison between accessible chromatin profiles using cosine similarity metric. Our data indicated a high degree of agreement of cis-regulatory element distributions between our PM_Lum samples (Cosine similarity *μ*_*Sc*_=0.82 ± 0.07; Cosine similarity) (**Figure 1b**). To identify whether our catalogue of accessible cis-regulatory elements was representative of other ER+PR+ breast tumors, we leveraged TCGA ATAC-seq data derived from bulk ER+PR+ tumor tissues (n=41; TCGA_Lum)^34^. Compared to our cohort, TCGA_Lum showed a higher number of unique accessible cis-regulatory elements (272,291 peaks; 289.89Mb) that encompassed 93.6% (93,172/99,516 peaks) of our PM_Lum catalogue. Of note, the PM_Lumour accessible cis-regulatory elements represented only 34.2% of the TCGA_Lum catalogue (**Figure 1c**), suggesting our depletion of immune cells may have enhanced the signal specific to cancer cells. Consistent with this observation, we estimated that our analysis led to the mapping of 88% of accessible chromatin within our cohort of 26 samples while the TCGA_Lum cohort reached similar saturation (87%) with 41 samples (**Figure 1d**). Thus, we established a catalogue from our PM_Lum cohort of high-confident accessible chromatin regions that were found across our cohort and illustrate a high level of robustness by being found almost entirely within the independent TCGA_Lum catalogue.

We next characterized the genomic distribution of accessible cis-regulatory elements across different genomic features (e.g. promoters, distal regions, coding exons UTRs, and intronic regions) within the PM_Lum and TCGA_Lum catalogues. Using the CEAS tool to estimate the relative enrichment level of accessible regions in gene features ^35^, we found that on average 36% of cis-regulatory elements mapped to promoters, 23% to introns, 23% to intergenic regions, 14% to UTR and 3% coding exons (**Figure 1e**). Using this same approach, we found that the cis-regulatory elements captured in the ATAC-seq data from the TCGA_Lum cohort had a similar distribution of intergenic regions (PM_Lum=23%, TCGA_Lum=26%; p=0.10) and coding exons (PM_Lum=3%, TCGA_Lum=3%; p=0.42). However, in contrast to the PM_Lum cohort the TCGA_Lum shows a higher distribution to introns (TCGA_Lum=39%, PM_Lum=23%; p<0.001) and a lower distribution to promoters (TCGA_Lum=26%, PM_Lum=36%; p<0.001) and UTRs (TCGA_Lum=6%, PM_Lum=14%; p<0.001) (**Figure 1e**). Thus, our results highlight that both PM_Lum and TCGA_Lum accessible chromatin catalogues favor non-coding regions, where most accessible regions are found in the promoter, intergenic and intronic sequences as opposed to the coding exons.

Considering that we used cell sorting to exclude immune cells from our tumor samples using an anti-CD45 antibody, we examined whether the difference in accessible chromatin profiles that we saw between TCGA_Lum and PM_Lum was due to immune infiltration. We tested for this immune infiltrate by comparing the similarity of the PM_Lum and TCGA_Lum profiles to a known immune reference comprised of publicly available chromatin accessibility data (DNaseI) from 12 immune cell types (trophoblast, CD1c+, myeloid progenitors, CD14+ monocytes, T helper17, T helper1, T helper2, CD8+alpha T cells, naive thymocytes T cells, CD4+ alpha-beta T cells, natural killer cells and B cells). Our results showed that the TCGA_Lum chromatin accessibility profile was significantly more similar to the accessible chromatin profile for 9 of the 12 immune cell types (trophoblast, CD1c+, myeloid progenitors, CD14+ monocytes, T helper17, T helper1, T helper2, CD8+alpha T cells, naive thymocytes T cells) compared to the PM_Lum profile (**Figure 1f**; *P* < 0.001, one-sided t-test). The accessible chromatin profile for 3 of the 12 immune cells tested (CD4+ alpha-beta T cells, natural killer and B cells) were not significant given the fact that CD45RO is not expressed in CD4+ T cells, natural killers and B cells ^36^. Altogether, our data suggests that although there are similarities between TCGA_Lum and PM_Lum, the cell sorting performed on our PM_Lum cohort led to a depletion of immune cells, resulting in a more cancer-cell-specific accessible chromatin catalogue.

Cis-regulatory elements work through the recruitment of transcription factors that bind to unique DNA recognition sequences. We therefore assessed the sequence composition of cis-regulatory elements from ER+PR+ breast tumors through DNA recognition motif enrichment analysis. Using the JASPAR database as a reference for motif recognition sites and the pan-cancer ENCODE DNase I hypersensitive sites as a background, we utilized the CentriMo method to identify 40 significantly enriched DNA recognition motif families, 6 of which are known to play an important role in luminal breast cancers: AP-2 (TFAP2A), Forkhead (FOXA1), STAT (STAT3), C/EBP (CREBBP), NR1 (RORA), GATA (GATA3) ^22,37–39^ (*P* < 0.001; Fisher’s exact test) (**Figure 1g, Table S3.1**). To corroborate our findings, we performed a similar DNA recognition motif enrichment analysis on the TCGA_Lum catalogue. We identified 57 DNA recognition motif families enriched in this cohort; 33/57 overlapped with the motifs enriched in our PM_Lum catalogue, 24/57 were unique to the TCGA catalogue, and 7/40 (HSF, MyoD/ASC, RHR, MADS, NFAT, NF and, B-ATF) were found only in the PM-Lum catalogue (**Figure S1**; **Table S3.2**). Some of which have been linked to breast cancer development and drug resistance ^37,40–43^. Together, these results demonstrated that our PM_Lum catalogue defines a broad spectrum of motif recognition sites, 82.5% of which are also found in the TCGA_Lum catalogue and 6 which are established markers of luminal breast cancer biology, thus reflecting the luminal breast cancer specificity of our catalogue.

### Individual cis-regulatory elements are rarely recurrently mutated

The enrichment of mutations within promoters and enhancers of key breast cancer genes, such as *TERT*^*14,27*^ and *FOXA1*^*25*^, suggests potential for recurrent mutations in additional regulatory regions. To search for other mutations in cis-regulatory elements in ER+PR+ breast cancer, we integrated our PM_Lum catalogue with somatic mutations from 348 ER+PR+ breast cancers in two whole-genome sequencing (WGS) breast studies (ICGC-EU^1^; n=306 and ICGC-US^44^; n=42). Of the 1,048,537 mutations found across whole-genome sequencing of ER+PR+ breast cancer samples from ICGC-EU and ICGC-US, an average of 1.7% (ICGC-US=1.76%; ICGC-EU=1.78%) [0.7%-3.4%; *n_SNVs*: min=4,295, max=35,650] were detected within our PM_Lum catalogue, which comprises 1.3% of the genome (**Figure 2a**). To identify whether our PM_Lum catalogue captured mutations specific to ER+PR+ breast cancers, we compared the localization of mutations to 19 ICGC WGS cancer cohorts **(Table S4)**. We found that these additional 19 cancer types all had significantly lower fractions of mutations overlapping our PM_Lum catalogue as compared to ER+PR+ breast cancer samples, with the exception of BOCA, PAEN-AU, and PRAD-UK (**Figure 2a)**. We then performed the same analysis using the TCGA_Lum catalogue of accessible chromatin and found similar results, with luminal breast tissue having a higher percentage of mutations localized to this region when compared to other tissues (p<0.01, two sided t-test; **Figure S2a**). These results highlight that mutations with luminal breast cancers are predominantly found within our accessible chromatin catalogue, thus setting the stage for interpreting mutations in cis-regulatory elements relevant to luminal breast cancer biology.

**Figure 2:**
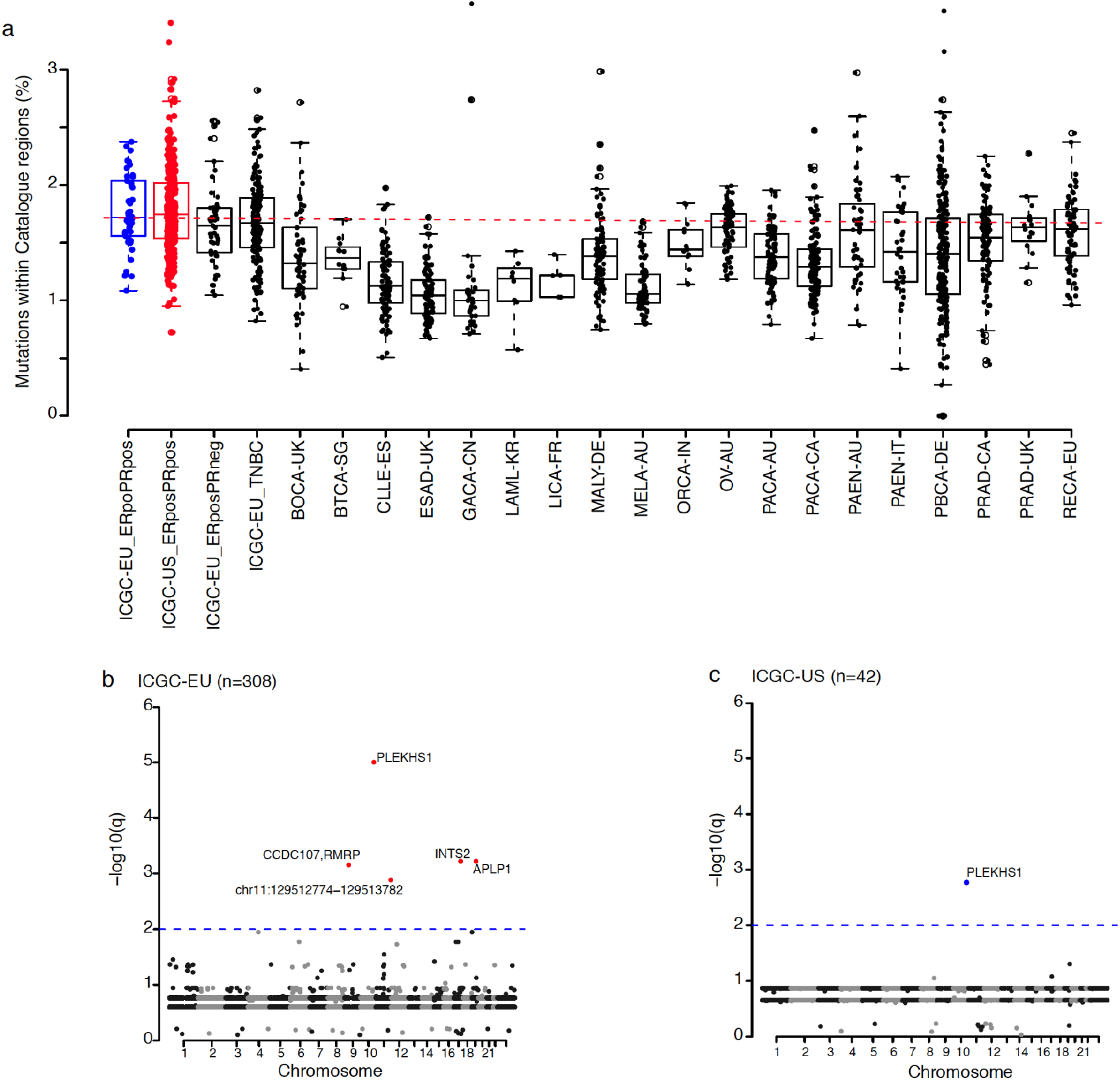
Mutation enrichment at cis-regulatory elements in ER+PR+ breast cancer. (**a**) Boxplot showing the percentage of regions from PM_Lum catalogue overlapping mutation calls from WGS from multiple cancer types. (**b**,**c**) Manhattan plots indicating regulatory regions significantly enriched in mutations using our in-house algorithm. The PM_Lum catalogue was used as accessible chromatin targets and the ICGC_EU WGS (**b**) or ICGC_US (**c**) was used as mutation calls. Dotted lines indicate q =< 0.01.

To identify highly mutated regulatory elements in ER+PR+ breast cancer, we analyzed frequently mutated regulatory elements using the ActiveDriverWGS method^19^. Restricting our analysis to our PM_Lum catalogue as the target regions, we found no driver mutations after multiple testing correction using two separate data sets, ICGC-EU (**Figure S2b, Table S5.1**) and ICGC-US (**Figure S2c, Table S5.2**) WGS data (q < 0.01; FDR). By running a similar analysis on the TCGA_Lum catalogue, ActiveDriverWGS identified one highly mutated distal region (chr10:8115662-8116163) using ICGC-EU (**Figure S2d**) and none using ICGC-US (**Figure S2e**). Although ActiveDriverWGS is a robust tool for calling drivers in regulatory elements, it takes a conservative one-to-one approach between mutations and active elements, negating the cumulative effect of multiple mutations within a hotspot region. To address these limitations, we designed an algorithm (HoRSE; Hotspot of cis-Regulatory, Significantly-mutated Elements) that relaxes the stringency of ActiveDriverWGS by looking for clusters of hotspot mutations within regulatory elements against a background of global and local somatic mutation rates (**Figure S2f;** Online methods). Using HoRSE, we found 5 unique cis-regulatory elements enriched for somatic mutations across ICGC-EU and -US (n_ICGC-EU_=5, n_ICGC_US_=1; q < 0.01, exact binomial test) with *PLEKHS1* being the only cis-regulatory element significantly enriched in both WGS cohorts (ICGC-EU: n=12/308; ICGC-US: n=6/42). (**Figure 2b**,**c, Table S5.4**). Two of the somatic mutations identified within the *PLEKHS1* promoter are thought to be attributed to APOBEC DNA-editing activity ^1,45^. In the ICGC-EU dataset, we identified 4 cis-regulatory elements enriched for somatic mutation in addition to *PLEKHS1* (Promoters of *INTS2;* n=6/308, *APLP1;* n=6/308, *and CCDC107/RMRP*^*25,45*^; n=6/308, and *Distal Region:* chr11:129512774-129513782; n=7/308) (**Figure 2b**,**c, Table S5.3**). Additionally, by applying our algorithm on the regions covered by the TCGA_Lum catalogue, we revealed 19 significantly mutated regions in the ICGC-EU dataset regions including CCDC107/RMRP (**Figure S2g**) and 3 regions in ICGC-US (promoter: RARA and 2 distal regions: chr8:98131092-98131993 and chr17:38603438-3860433; **Figure S2h**). Our results highlight the small number of recurrent mutational hotspots across all the cis-regulatory elements of luminal breast cancer. Thus, similar to how the search for driver genes is hindered when focusing on single exons, our results show the hunt for cancer drivers within individual cis-regulatory elements may be too limiting resulting in the few observed recurrently mutated regions.

### Non-coding mutations reveal cancer driver cistromes in luminal breast cancer

The genome can be looked at as a collection of cis-regulatory elements that can be organized into cistromes, based either on the DNA recognition sequence content or on actual occupancy by transcription factors. As our previous analysis highlights the limitations of identifying drivers using individual cis-regulatory elements, our next step was to assess the presence of cancer driver cistromes in ER+PR+ luminal breast tumors. First, we measured the enrichment for DNA recognition motifs within the PM_Lum catalogue of cis-regulatory elements found to be mutated in primary luminal breast tumors from the ICGC-EU and -US studies. This revealed significant enrichment for several DNA recognition motifs related to the JUN, FOS, Forkhead, NFAT, POU and REL families of transcription factors across both ICGC dataset (**Figure 3a**). The NF1, C2H2, IRF and HD-CUT DNA recognition motifs were uniquely enriched in cis-regulatory elements mutated based on the ICGC-EU dataset (**Figure 3a**).

**Figure 3:**
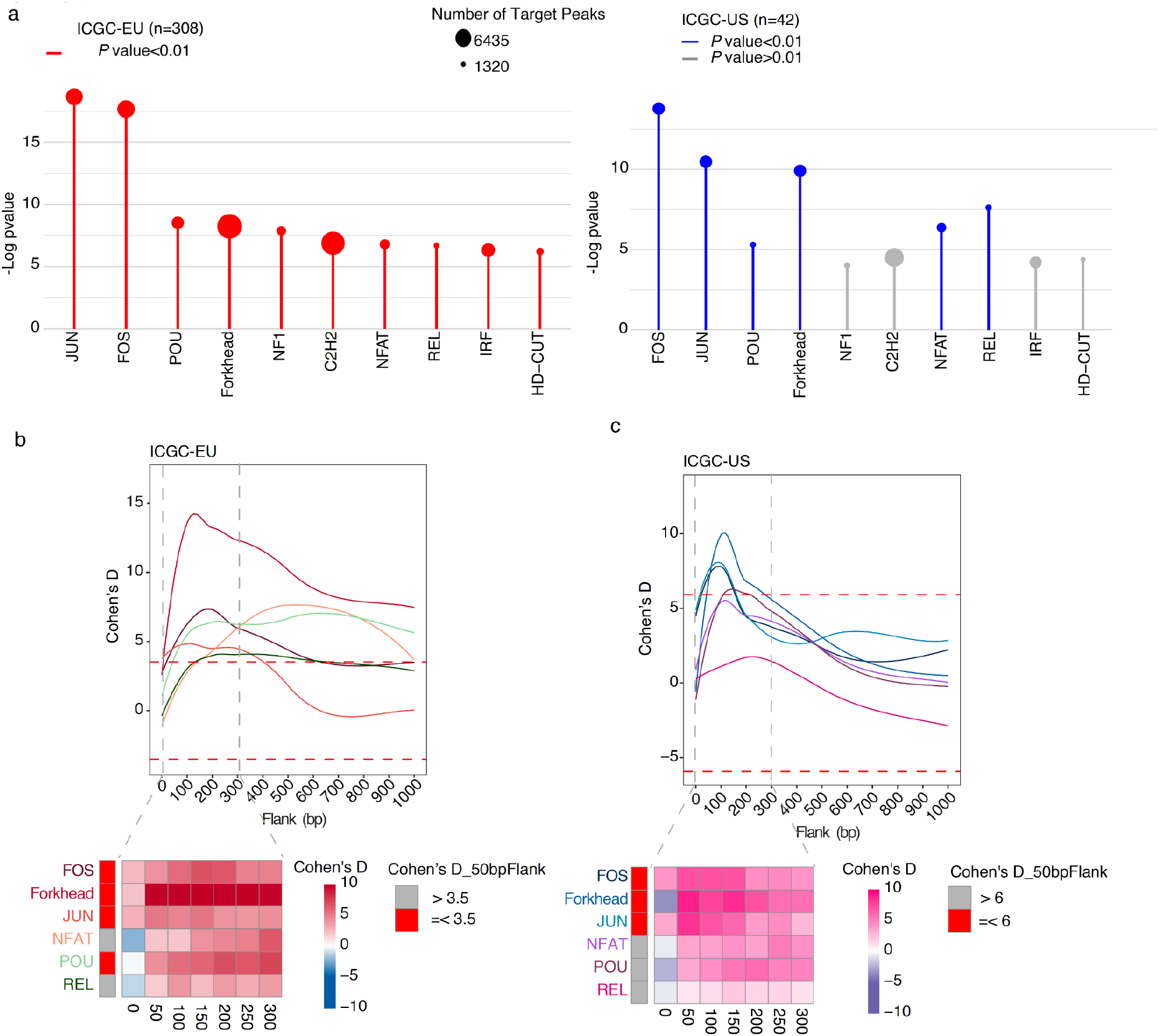
Mutation analysis at recognition sites of motifs enriched in ER+PR+ breast cancer. (**a**) Lollipop graph showing enriched motif families in PM_Lum catalogue overlapping SNVs from ICGC-EU (Red) and ICGC-US (Blue) against the total PM_Lum catalogue (p-value < 0.01; Grey: p-value > 0.01). (**b**,**c**) graph (Up) and heatmaps (Bottom) showing the enrichment of mutations at DNA recognition sites found to be significantly enriched in the PM_Lum catalogue using ICGC-EU **(b)** and ICGC-US **(c)** mutation calls. Cohen’s D was calculated based on resampling and the value indicates significant enrichment. The red dotted line indicates Cohen’s D median.

To focus on DNA recognition motif-based cistromes relevant to luminal breast cancer, we subdivided cis-regulatory elements from our catalogue of accessible chromatin regions based on the presence of DNA recognition motifs enriched in mutated cis-regulatory elements across both ICGC-EU and ICGC-US datasets, namely JUN, FOS, Forkhead, NFAT, POU or REL. We calculated the frequency of mutations across varying window sizes (0 to 1,000bp) around the cis-regulatory elements from each of the motif-based cistromes using modMEMOS (modified Mutation Enrichment within the Motifs and Flanking Regions; **Figure S3;** Online methods)^17,25^. We estimate the effect size of mutation enrichment in DNA recognition motifs compared to a background model using Cohens’ D, a statistical value that represents the standardised difference between two means. Using a window of 50 bp flanking the motif recognition sites, as defined by the work from Mazrooei *et al*. ^17,25^, we found an enrichment for mutations near the JUN, FOS and Forkhead motif-based cistromes in both ICGC-EU (**Figure 3b**) and ICGC-US data sets (**Figure 3c**). Additionally, cis-regulatory elements proximal to POU motif cistrome were found to be enriched in mutations uniquely in the ICGC-EU dataset (**Figure 3b)**. These results suggest that non-coding mutations preferentially accumulate across cis-regulatory elements that harbor specific DNA recognition motifs, namely JUN, FOS or Forkhead motifs.

Given that transcription factors of the same family can bind the same DNA recognition motif, we explored the transcription factor-based cistromes to examine whether these variants are targeting transcription factor binding sites specific to breast tumors. We leveraged the publically available collection of ChIP-seq datasets of transcription factors (n=48) and co-factors (n=30) from the luminal breast cancer cell line (MCF7) ^46^to identify luminal specific transcription factor-based cistromes. We first clustered all cistromes according to their similarity in ChIP-seq signal across our catalogue of cis-regulatory elements from luminal breast tumors and identified 7 distinct clusters (**Figure S4**), including one consisting of the ESR1, FOXA1 and GATA3 transcription factors (TFs_1). We next used modMEMOS to quantify the enrichment of mutations over these cistromes. Using the mutation calls from the ICGC-EU dataset, we identified 28 cancer driver cistromes (AHR, AR, CEBPB, CREBBP, CTCF, ELF1, ESR1, FOSL2, FOXA1, FOXM1, GABPA, GATA3, JUND, MAX, MYC, NR2F2, REST, TCF12, TEAD4, TFAP2A, TFAP2C, and ZNF217) (**Figure 4a**). We further refine these transcription factor-based cistromes by including only the cis-regulatory elements that harbor a matched DNA recognition site for the designated transcription factor family. Using modMEMOs on these DNA recognition site-specific transcription factor-based cistromes, we identified 10 cancer driver cistromes (CTCF, ELF1, ESR1, FOSL2, FOXA1, FOXM1 GATA3, JUND, TFAP2A, and TFAP2C) that are enriched in mutations in both the ICGC-EU (**Figure 4b**) and ICGC-US (**Figure 4c**) datasets. Consistent with the motif-based cistromes that we identified as cancer drivers (**Figure 3b**,**c**), we observed a similar enrichment of most, but not all transcription factor-based cistromes that compose each motif family (Forkhead, JUN and FOS), with the exception of the REL motif family (**Figure S5**). Furthermore, we found that in the majority of cases, not all transcription factor-based cistromes for a given motif family defined cancer driver cistromes (e.g. Forkhead motif family). Rather mutations were found to be enriched in specific transcription factor-based cistromes (**Figure S5**). Altogether, our results highlight that key transcription factor-based cistromes are cancer drivers independent of their motif families or based on their similarity to other cistromes, indicating the mutations are selectively enriched within specific driver cistromes.

**Figure 4:**
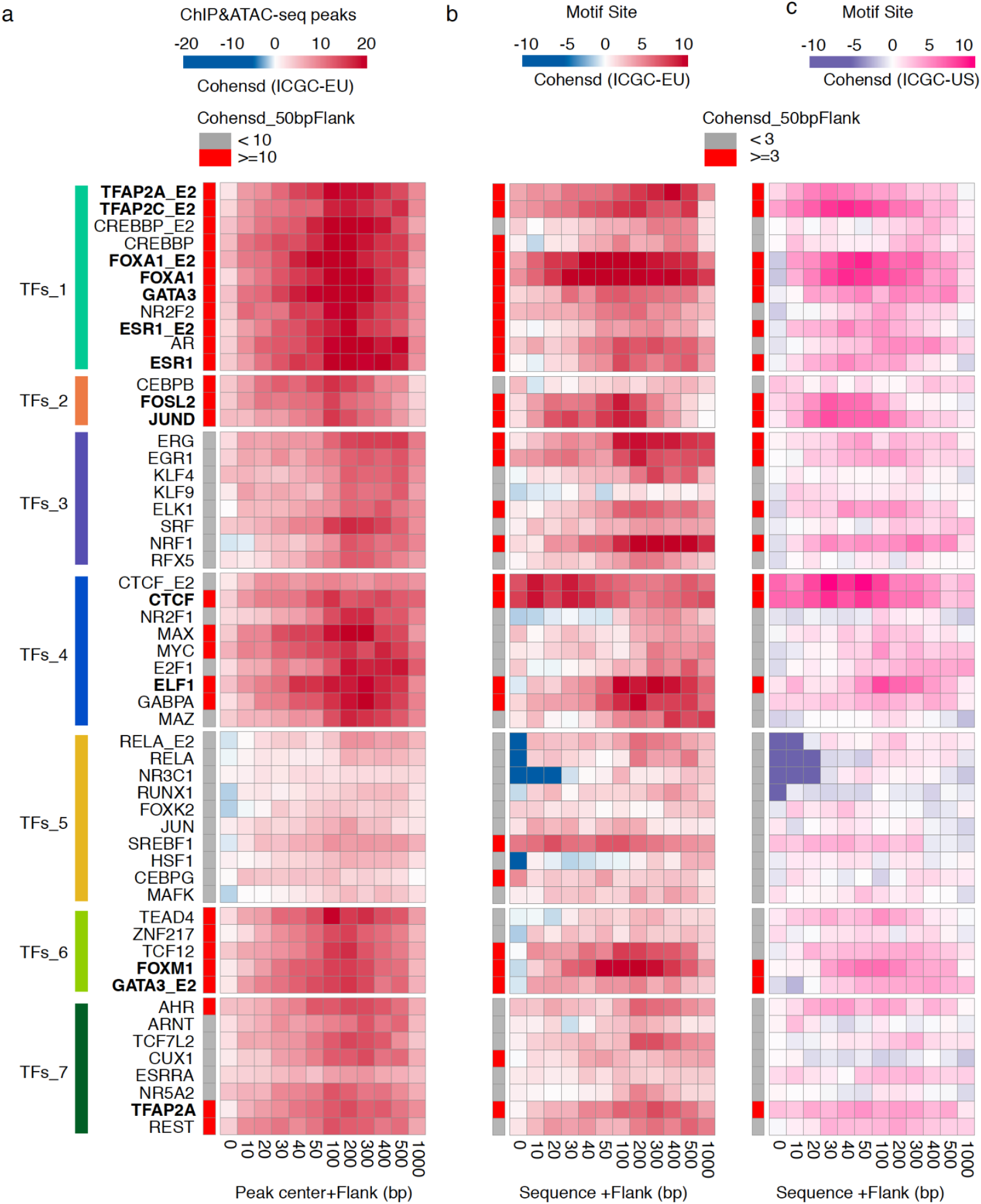
High enrichment of mutations at cistromes of key transcription factors involved in ER+PR+ breast cancer. Heatmaps showing enrichment of mutations at ChIP-seq peak centers and flanking regions (0-1000bp) using ICGC-EU WGS dataset (**a**), transcription factor binding sets using ICGC-EU (**b**), and ICGC-US WGS datasets (**c**). Cohen’s D was calculated based on resampling and the value indicates significant enrichment (Enrichment > Median (Cohen’s D)).

We next examined if the non-coding mutations within or flanking (100bp) the DNA recognition motif found within the cancer driver cistrome for CTCF, TFAP2C, GATA3, FOXA1, ESR1, FOSL2, JUND, TFAP2A, ELF1, and FOXM1 could alter transcription factor binding to the chromatin. Using the intragenomic replicate (IGR) method^22^ predicted that less than 40% of the non-coding mutations could alter the binding intensity of any of these transcription factors to the chromatin (CTCF: Down=36%, Up=15%; TFAP2C: 31%,28%; GATA3: 19%,10%; FOXA1: 14%,17%; ESR1: 18%,9%; FOSL2: 27%,13%; and JUND: 14%,5%; TFAP2A: 32%,20%; ELF1: 33%,18%; FOXM1:20%,13%) (**Figure S6**). These results argue that despite the enrichment of mutations observed over transcription factor-based cistromes, only a minority of these mutations can directly impact the binding affinity of transcription factors to cis-regulatory elements.

### Cancer driver cistromes correspond to transcription factors essential to luminal breast cancer

To better understand why specific transcription factor-based cistromes are enriched for non-coding mutations in luminal breast cancer, we examined whether this enrichment reflected the dependency to some as opposed to all transcription factors expressed in luminal breast cancer. Using the genome-wide shRNA essentiality screen data from luminal breast cancer cell lines generated as part of the DepMap project ^47,48^, we found that 4 of the 10 transcription factors linked to cancer driver transcription factor-cistromes were exclusively essential in luminal breast cancers (GATA3, ESR1, FOXA1, TFAP2A) and five additional transcription factors were essential in all breast cancers, regardless of subtype (CTCF, FOXM1, TFAP2C, JUND and FOSL2) (**Figure 5, p < 0.05**). ELF1 was the only transcription factor linked to a cancer driver cistrome not essential in luminal breast cancer cells (**Figure 5**). While we found that the CREBBP and CEBPG transcription factors were essential preferentially in luminal breast cancer, we did not identify these transcription factor-cistromes as cancer drivers as they were only significantly enriched in mutations in the ICGC-EU dataset. Altogether these results support the identification of cancer driver cistromes based on transcription factors that are essential to the growth of luminal breast cancer.

**Figure 5:**
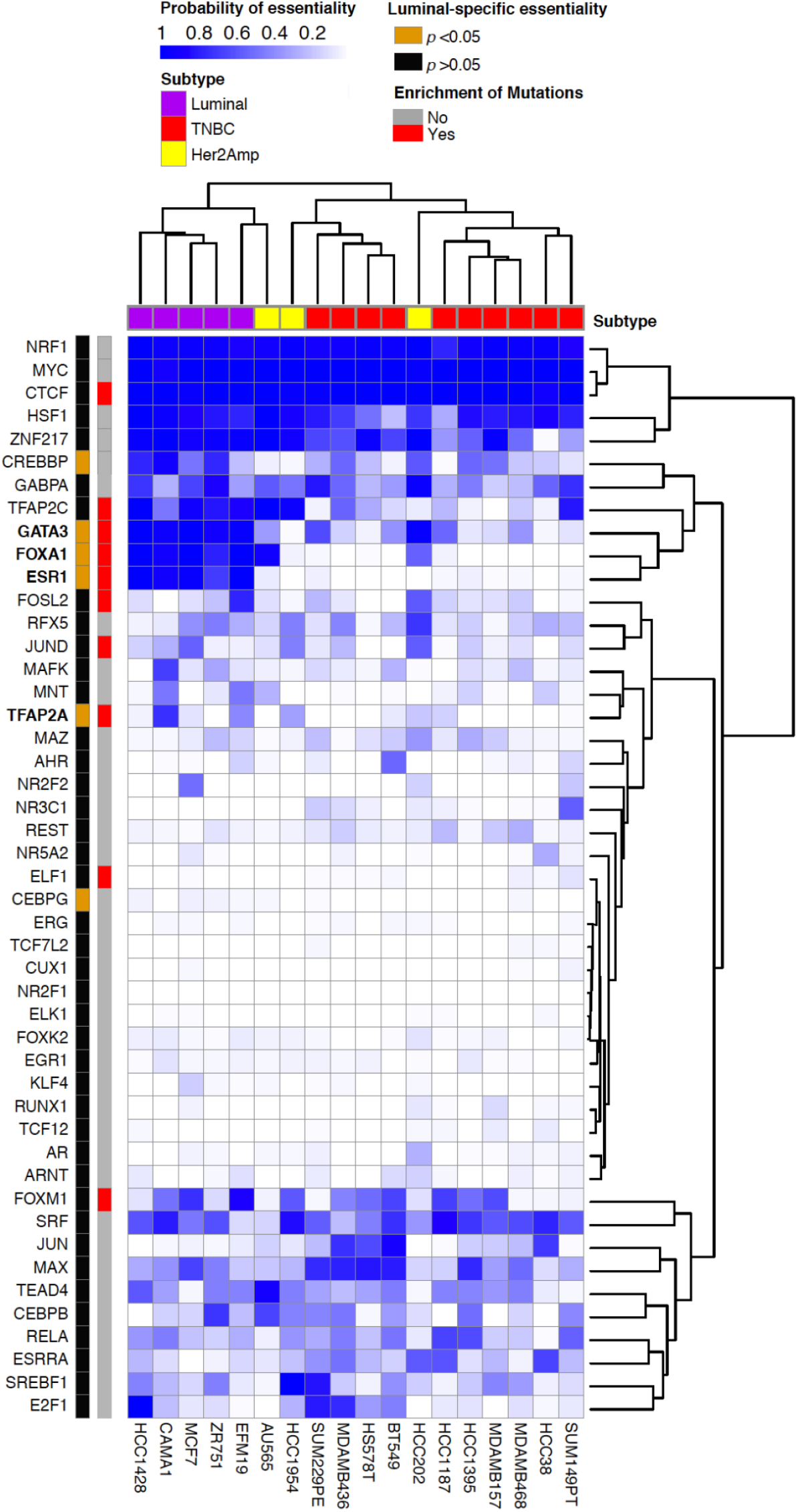
Cancer driver cistromes are of transcription factors essential to luminal breast tumors. A heatmap showing the probability of the essentiality of the transcription factor in several breast cancer cell lines with different subtypes (Luminal, TNBCs, and HER2). Column annotation indicates the enrichment of mutations at binding sites +/- 50bp, and rows annotation shows cell line subtype.

## Discussion

Our study depicts the cancer driver cistromes specific to luminal ER+PR+ breast cancers as identified by an enrichment of non-coding mutations flanking DNA recognition motifs of cis-regulatory elements accessible in luminal breast tumors. Using flow-sorting to enrich the cancer cell population, we generated a robust catalogue of luminal-specific accessible chromatin regions. Within this catalogue, we identified seven recurrently mutated cis-regulatory elements that occur at a low frequency. By expanding our search to transcription factor-based cistromes, we identified 10 cancer drivers and showed that a minority of the non-coding mutations can directly impact the transcription factor binding to cis-regulatory elements. Finally, we show these 9 out of the 10 transcription factor-cistromes are essential to breast cancer, and 4 of which are specific to luminal breast cancer.

Somatic variants and genomic rearrangements affecting the protein-coding regions of luminal breast cancers have been well-characterized ^1,2,49,50^, these regions account for less than 2% of the genome^51,52^. The importance of acquired genetic variants found in cis-regulatory elements is highlighted in a luminal breast cancer study by Bailey et al. ^3^ and across multiple breast cancer subtypes by Rheinbay et al. ^25^. Bailey et al. identified several somatic mutations with functional consequences within the promoters and enhancers that regulate the *ESR1* gene ^3^. The study by Rheinbay et al. describes somatic mutations across several promoters, including *FOXA1*, and their effect on gene expression ^25^. Our analysis of the mutation burden within luminal ER+PR+ breast cancer cis-regulatory elements yielded only seven significant hits. Across both the ICGC-US and -EU cohorts, we found significant enrichment of mutations in the *PLEKHS1* promoter that is likely a result of APOBEC DNA-editing activity ^1^, however, this region is also known as a genetic marker of aggressiveness for differentiated thyroid carcinomas ^53^. Although significant, our results show that the hunt for cancer drivers within individual cis-regulatory elements is limiting at best, resulting in the few observed recurrently mutated individual cis-regulatory elements. Discovering cancer driver mutations in the non-coding space is challenging due to heterogeneity in the cis-regulatory element and mutational space between individual tumors, leading to a need of large datasets to identify rarely occurring cancer driver mutations ^52^.

As individual cis-regulatory elements are functional units of the cistrome, akin to how exons make up a gene, we expanded our search for cancer drivers by partitioning our accessible chromatin region into cistromes specific for luuminal breast cancer. GWAS studies have identified thousands of risk variants linked to diseases including breast cancers ^3,17,22,25,54^. In luminal breast cancer a number of these risk variants have been shown to accumulate at the cistromes of key transcription factors in luminal breast cancer, namely ESR1 and FOXA1 ^3,22,55^. The CTCF/cohesin binding sites, regulators of the 3D structure of chromatin, are enriched in point mutations in a highly stereotypic pattern across various cancer types which may affect transcriptional regulation and result in genomic instability ^56^. Additionally, Mazrooei et al. showed enrichment of mutations within the cistrome of master regulators of prostate cancer such as FOXA1, HOXB13 and AR^17^. Our study provides a look into another aspect of cancer driver search by looking at mutation load within motif and transcription factor-based cistromes. We detected an enrichment of mutation at regions flanking the DNA recognition motif in cistromes crucial to luminal breast cancer, namely the cistromes of CTCF, TFAP2C, GATA3, FOXA1, ESR1, FOSL2, JUND, TFAP2A, ELF1, and FOXM1. The biological significance of mutagenic processes occurring at the flanking regions of cistrome over the active binding sites is yet to be fully understood but is a phenomenon seen in prostate cancer ^17^. While other studies in melanoma ^57,58^, lung ^59^, and colorectal ^56^ cancers have found the inverse true, they have attributed this mutational enrichment to restricted DNA-accessibility affecting repair machinery due to either chromatin conformation change, or occupancy of specific transcription binding sites by proteins ^60^. Approximately 5-36% of these mutations are predicted to impact transcription factor binding to the chromatin. Altogether, we describe an increase of mutational burden at specific cistromes defining them as cancer driver cistromes.

As validation of our cancer driver cistromes, we determined from the DepMap project ^47,48^ that four transcription factors associated with our driver cistromes were preferentially essential to luminal breast cancers: GATA3, ESR1, FOXA1 and TFAP2A. Among those, GATA3, ESR1 and FOXA1 have been widely shown to be involved in luminal breast cancer development and resistance to endocrine therapy^61^, while TFAP2A is associated with the luminal breast phenotype ^39^. Five additional transcription factors, CTCF, FOXM1, transcription factor AP2C, JUND and FOSL2, were essential across all breast cancer cell lines. While not luminal exclusive, these transcription factors have roles in breast cancer progression, aggressiveness, cell motility, modulating cancer cell proliferation, and response to therapy ^39,62–66^. In conclusion, our study provides new insights to identifying cancer drivers beyond the protein-coding space to benefit the development of precision medicine from cancer driver events applicable to breast and other cancer types.

## Material and methods

### Patient tumor samples

Twenty-six primary tumors were obtained from surgical specimens of patients with ER+PR+ invasive ductal carcinoma. Patients’ consent and tumor stratification were obtained through UHN living biobank under REB # 16-5524.

### Tumor processing and ATAC-seq library preparation

Breast tumors were minced into small pieces and digested at 37C, in mammary Epicult (STEMCELL Technologies, Vancouver, BC, Canada) media supplemented with 10% FBS (WISENT, ST-BRUNO, QC, Canada) and collagenase (STEMCELL Technologies, Vancouver, BC, Canada), and further dissociated in 5 mg/ml dispase for 2min. Cells were counted and live cells sorted into two populations, immune and malignant cells enriched using sytox blue (ThermoFisher Scientific, Massachusetts, USA) and anti-CD45 antibody (ThermoFisher Scientific, Massachusetts, USA). Fifty thousand were used for ATAC-seq library preparation as described previously ^31^. Briefly, cells were lysed for 5 min followed by transposase reaction and library amplification using Nextera DNA Library Prep Kit (Illumina, California, USA). Libraries were then size-selected (240-360 bp) using PippinHT (Sage Science, Beverly, CA, USA) and sequenced (NextSeq 550) using 50 bp single reads.

### ATAC sequencing and data analysis

Reads were aligned to hg19 using bowtie2/2.0.5 using default parameters. Aligned reads were then filtered by removing duplicated and mitochondrial reads using samtools/0.1.18. We then used MACS2/2.0.10^33^ to call accessible chromatin peaks using the following parameters: macs2 callpeak -t {input.bam} -g hs --keep-dup all -n {sample-name} -B --nomodel –SPMR -q 0.005 --outdir {OutputDir}

### Enrichment of Genomic Features in Open Chromatin Regions

The open chromatin regions from ATAC-seq, represented using a BED file, were used as input for CEAS v1.0.2 ^35^ along with hg19 refGene, running the default ChIP Region Annotation and Gene-centered Annotation modules. Similarity between ATAC-profiles was estimated using all unique peaks in a pairwise-comparison between samples. A cosine similarity was used to negate the differences in global peak amplitudes and compare the relative amplitudes.

### Motif enrichment

We analyzed motif enrichment using CentriMo from the Meme-suite tool version 4.9.0_4 and as a reference, we used the JASPAR_CORE_2016.meme database. This analysis was run on multiple catalogues. First, we run PM-Lum and TCGA_Lum catalogues using as a background a catalogue of publicly available DNaseI sensitive sites identified in several cell lines. The DNaseI sensitive sites were downloaded from the Encyclopedia of DNA Elements (ENCODE). Next, we ran the same analysis on PM_Lum accessible chromatin regions that overlapped mutations from ICGC_EU and ICGC_US datasets using as a background the full PM_Lum Catalogue.

### HoRSE (Hotspot of cis-Regulatory, Significantly-mutated Elements)

In order to identify mutation enrichment within non-coding regions, we developed an algorithm that uses an exact binomial test for each region of interest against a sample-wide noncoding background mutation rate (https://github.com/pughlab/BCa_ATACSEQ_Project/tree/main/HoRSE). We first define the search space as the overlap between cis-regulatory elements and the ATAC-catalogue, as well as separate variants into non-coding and coding based on the UCSC hg19 knownGene annotations. By tiling a 5kb window across the cis-regulatory elements for the search space, we fit the number of variants found within the tiled cis-regulatory elements to a poisson model to estimate the average background mutation rate. We also used a 5kb sliding window approach to identify the loci within each cis-regulatory element with the highest mutation burden. The highest mutation burdens were compared to the background mutation rate using an exact binomial test and corrected for multiple hypothesis testing using an FDR correction.

### Mutation Enrichment at Motif sites (ModMEMOS)

To analyze the enrichment of mutations at motif sites, we used a modified version of the previously published tool MEMOS (ModMEMOS) (https://github.com/pughlab/BCa_ATACSEQ_Project/tree/main/modMEMOS) ^17^ (**Figure S3**). First, similar to the previous version, we scanned for motif sites using either the PM_Lum ATAC-seq Catalogue or PM_Lum ATAC-seq Catalogue that overlap publicly available ChIP-seq data run on MCF7 using MOODS/1.9.2 tool ^67^. The previously published version of MEMOS assumed a normal distribution of number of mutations, however, due to the low number of mutations within cis-regulatory elements, we adopted a poisson distribution to better fit our data. Additionally, MEMOS established the null distribution of mutation enrichment by randomly sampling from the entire genome followed by adding a flanking region, resulting in the potential for the background region to include the target regions. We address this by adding the maximum flanks (1000bp) to all motif recognition sites first, and then restricting sampling to all regions that do not overlap the ENCODE blacklist regions as well as all original motifs +/- 1000bps. Finally, MEMOS estimates its p-value for motif enrichment by calculating the distance of the number of mutations within the target cistromes from the standardized mean of the null distribution. Due to the low number of mutations within some of our cistromes, we opted for a confidence interval approach by resampling the target and background regions, followed by calculating mutation enrichment within the resampled regions and estimating the effect size of enrichment using Cohen’s D. From a technical perspective of modMEMOS, we added a flanking region (0-1000bp) to Motif sites/ChIP peak centers using Bedtools slop and resampled the resulting bedfiles 100 times, taking 80% of the bedfiles each time. In parallel, we generated a background bedfile by randomly shuffling all of the motif sites +/- 1000bp while excluding the Motif sites/peak center +/- flanking region as well as the ENCODE blacklist regions. Similar to Motif sites/ChIP peak centers, the background bedfile was resampled 100 times, taking 80% of the regions each time. Taking into consideration our regions of interest and background file we identified the regions that overlapped mutations from ICGC-EU and US datasets, and counted the number of mutations for each transcription factor site and flanking region. Finally, we compared the mutation counts from the region of interest to the background and calculated Cohen’s D using the following equation: “Mean difference / pooled standard deviation”. We determined the enrichment threshold based on the Cohens’ D median.

### Intra-Genomic Replicates (IGR)

To predict the effect of SNVs on transcription factors binding affinity, we run the Intra-genomic replicates (IGR) tool ^22^. In summary, IGR uses ChIP-seq data of the transcription factor of interest to analyze the change in signal intensity in regions harboring SNVs compared to surrounding regions. Herein, we analyzed the binding affinity of the transcription factor that binds sites found to be enriched in mutation. Our regions of interest were the transcription factor binding sites +/- 100bp flanking regions. We used the ICGC-EU mutation dataset as the SNVs file.

### Essentiality screens

Project Achilles genome-wide shRNA essentiality screen data was downloaded from the DepMap portal, specifically the “Achilles” dataset ^47,48^. The analysis was focused on breast cancer cell lines that showed consistency in subtyping according to all three genesets PAM50, SCMOD2, and SCMGENE. The probability of essentiality was used as a score 1 being most essential and 0 non-essential.

### Identifying luminal-specific essentiality

Enrichment of essentiality for one breast cancer type compared to the rest was calculated using an approach inspired by GSEA ^68^. The probability of essentiality (*P*_*e*_) values were assigned a direction based on whether they were part of the cancer type of interest (COI; positive) or not (negative). A curve was fitted to the ordered *P*_*e*_ list and the area under the curve (AUC) was calculated. An exact p-value for each cancer-type was calculated using a permutation test (n_perm=1000) where the cancer type index was randomized and the AUCs recalculated. All p-values were corrected for multiple testing using FDR. The standardized AUC was calculated based on a min/max AUC range, where the min is defined as *P*_*e*_=-1 for all non-COIs and *P*_*e*_=0 for all COIs, while the max has *P*_*e*_=0 for all non-COIs and *P*_*e*_=1 for all COIs.

## Declarations

### Ethics approval and consent to participate

The University Health Network Ethics Board operates in compliance with the Tri-Council Policy Statement reviewed and approved this project REB #16-5524.

### Availability of data and material

The ATAC-seq raw data generated from our PM_Lum cohorts were uploaded to EGA (European Genome-Phenome Archive) under accession code: EGAS00001005235.

### Availability of codes

https://github.com/pughlab/BCa_ATACSEQ_Project

## Funding

This research was supported by a grant from Susan G. Komen®.TJP holds the Canada Research Chair in Translational Genomics and is supported by a Senior Investigator Award from the Ontario Institute for Cancer Research and the Gattuso-Slaight Personalized Cancer Medicine Fund at the Princess Margaret Cancer Centre. Infrastructure support was provided by the Princess Margaret Cancer Foundation; Canada Foundation for Innovation, Leaders Opportunity Fund, CFI 340 #32383; and Ontario Ministry of Research and Innovation, Ontario Research Fund Small Infrastructure Program (TJP). This work was also supported by the Canadian Institute for Health Research (CIHR: Funding Reference Number 136963, 158225, and 168933 to M.L.) and the Princess Margaret Cancer Foundation (M.L.). M.L. holds an Investigator Award from the Ontario Institute for Cancer Research and the Bernard and Francine Dorval Award for Excellence from the Canadian Cancer Society.

SE is supported by CIHR Banting Postdoctoral fellowship

Project Grant from the Canadian Institutes of Health Research (CIHR) to J.R. and Investigator Award to J.R. from the Ontario Institute for Cancer Research (OICR). Funding to OICR is provided by the Government of Ontario.

## Authors’ contributions

SE designed workflows, performed tissue processing and library preparations, analyzed and interpreted the genomic data and did all subsequent analysis.

RQ developed algorithms

JH, PM, PG, HZ,JB, JR assisted in bioinformatic analysis

CY assisted in tissue processing

IC and YH performed macrodissection, tissue-staining, DNA/RNA extraction.

LEO organized the transfer of patients data

DJC assisted with the research ethic application and transfer of tissues

SD provided all pathological reviews of the tissues used in this project.

ML, and TJP designed, organized and managed this project.

## Acknowledgements

We thank the staff of the Princess Margaret Genomics Centre (www.pmgenomics.ca, Troy Ketela, Julissa Tsao, Nick Khuu and Monika Sharma) and Bioinformatics Services (Carl Virtanen, Zhibin Lu, Jin Qun, and Natalie Stickle) for their expertise in generating the sequencing data used in this study.

## Notes

### Competing Interest Statement

The authors have declared no competing interest.

