## Supplemental Figures 1-6 for "Non-coding mutations reveal cancer driver cistromes in luminal breast cancer"

Figure S1

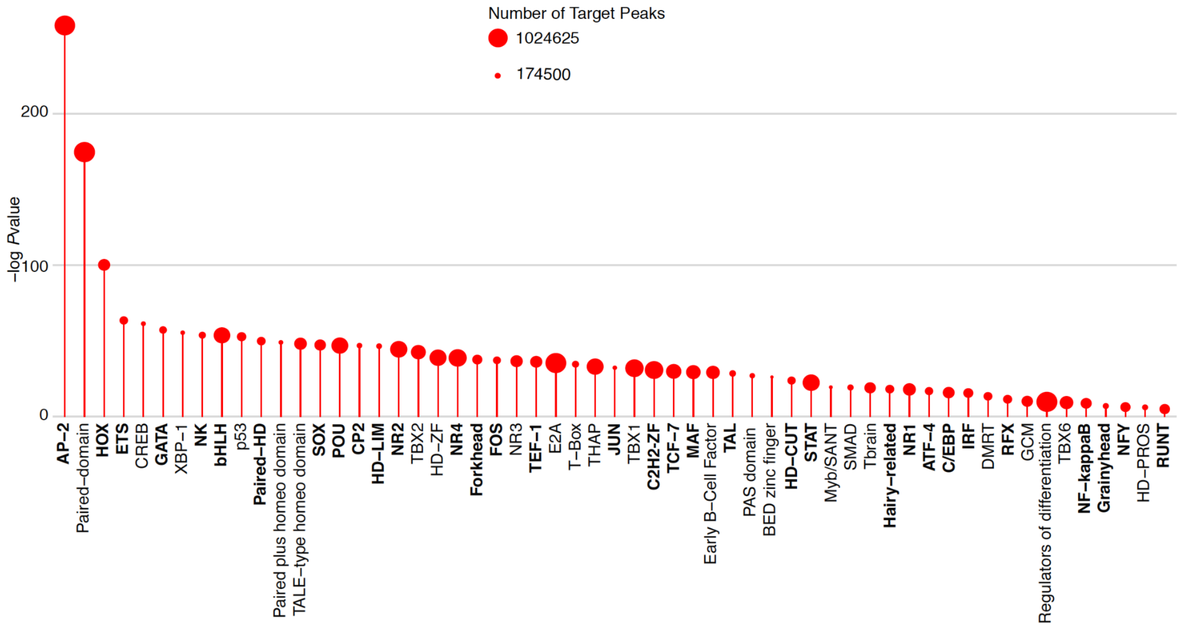


**Figure S1: Chromatin accessibility in ER+PR+ breast cancer from the TCGA dataset.** Lollipop graph showing enriched motif families in TCGA_Lum breast tumors (p-value < 0.01). The catalogue of 41 TCGA ATAC-seq data was used. Enrichment of motifs within ATAC-seq regions against DNaseI hypersensitive sites from several cell lines was computed. Motif families were obtained using Jaspar database.

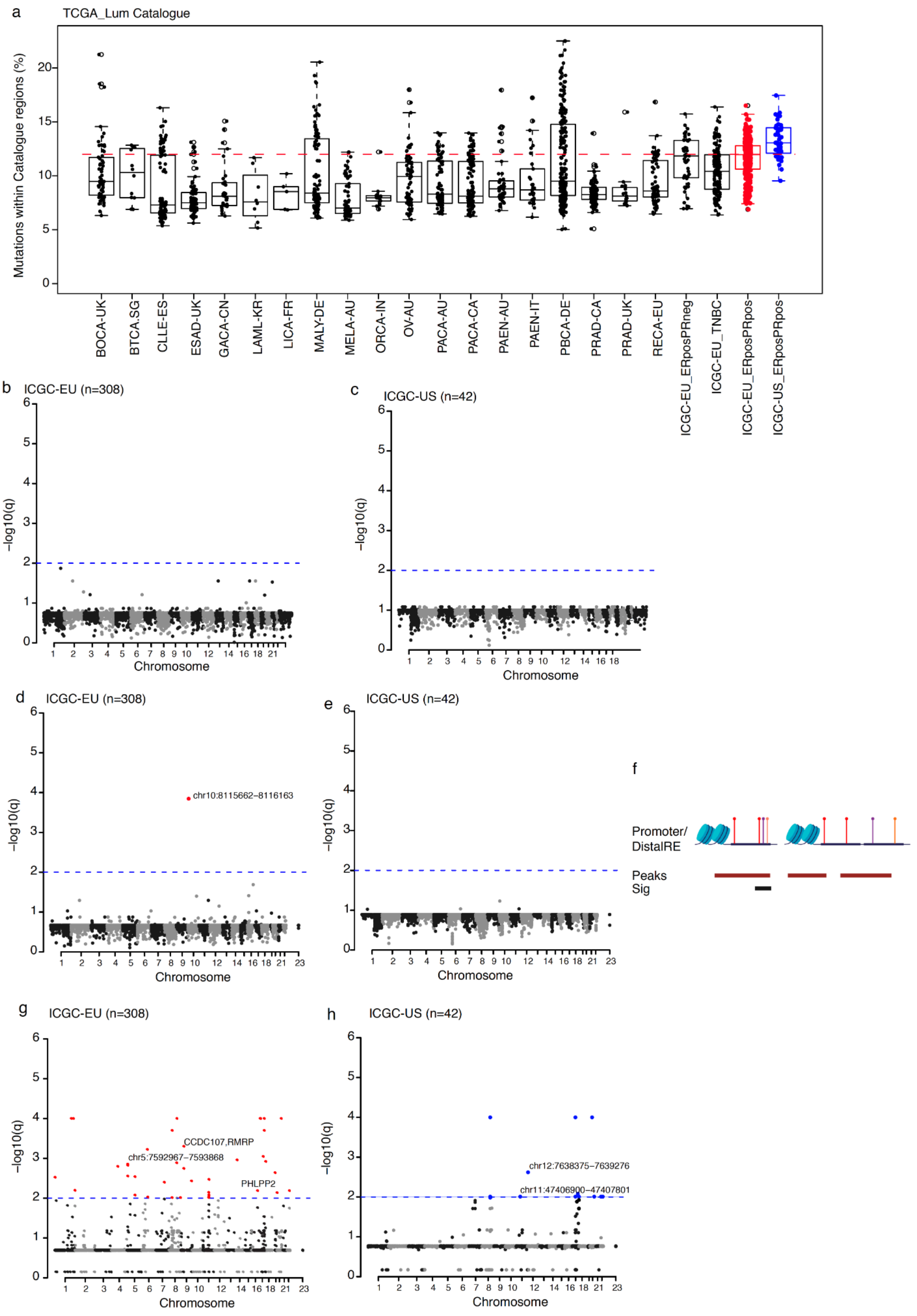


**Figure S2: Enrichment of mutations at individual cis-regulatory elements in the TCGA_Lum breast cancer cohort.**

(**a**) Boxplot showing the percentage of regions from TCGA_Lum catalogue overlapping mutation calls from WGS from multiple cancer types. (**b,c**) Manhattan plot showing

ActiveDriverWGS results using as the accessible chromatin target, our PM_Lum catalogue and as the mutation calls ICGC_EU WGS (**b**) or ICGC_US (**c**). (**d,e**) Manhattan plots showing ActiveDriverWGS results using as the accessible chromatin target, the publicly available TCGA_Lum catalogue and as mutation calls ICGC_EU WGS (**d**) or ICGC_US (**e**). (**f**) Schematic showing the CRE_Enrich algorithm. (**g,h**) Manhattan plots indicating regulatory regions significantly enriched in mutations using our in-house algorithm. The TCGA catalogue was used as accessible chromatin targets and as mutation calls we used the ICGC_EU WGS (**g**) or ICGC_US (**h**) datasets. Dotted lines indicate FDR =< 0.01.

Figure S3

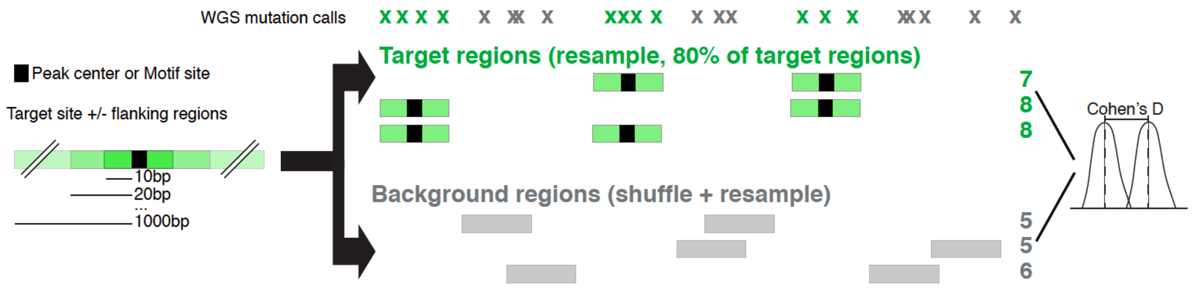


**Figure S3:** Schematic showing the adapted MEMOS analysis approach (Modified MEMOS; ModMEMOS).

Figure S4

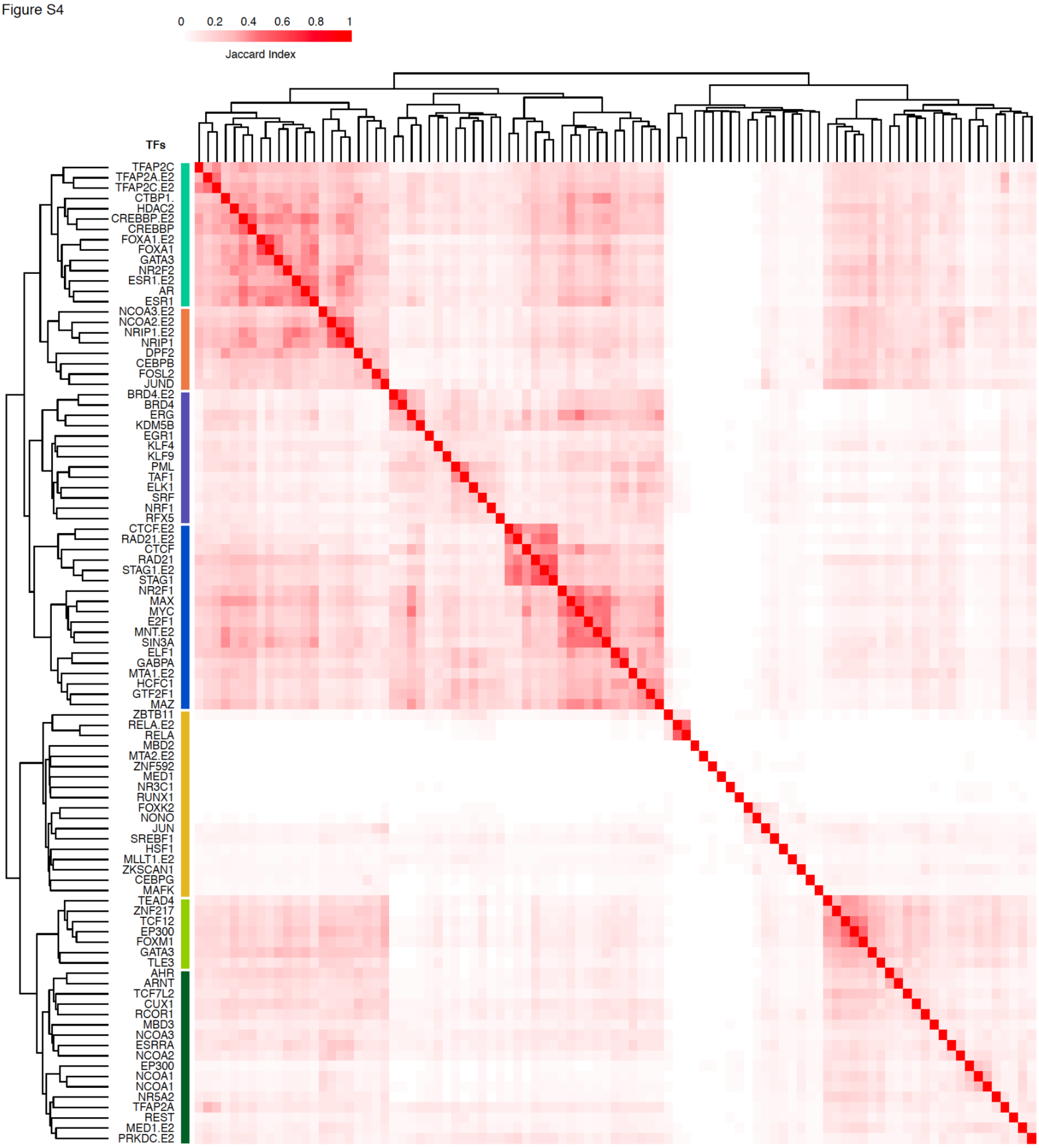


**Figure S4: Binding profile of transcription factors in MCFs.** Heatmap representing clustering of transcription factor ChIP-seq overlapping PM_Lum accessible chromatin in MCF7 using Jaccard index.

Figure S5

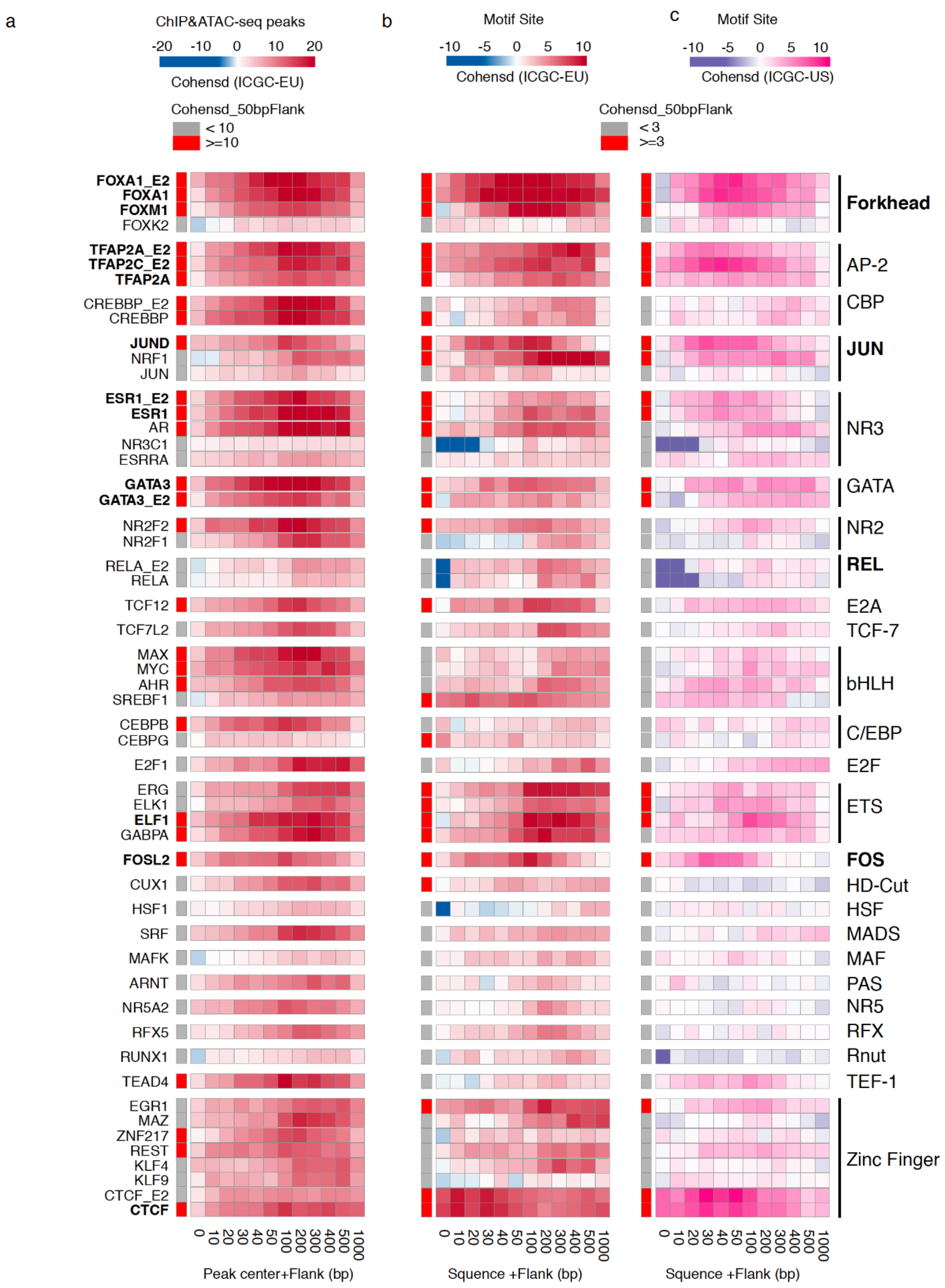


**Figure S5** | Heatmaps showing clustering of cancer driver cistromes based on motif families. Enrichment of mutations at ChIP-seq peak centers and flanking regions (0-1000bp) using ICGC-EU WGS dataset (**a**), transcription factor binding sets using ICGC-EU (**b**), and ICGC-US WGS datasets (**c**). Cohen's D was calculated based on resampling and the value indicates significant enrichment (Enrichment > Median (Cohen's D)).

Figure S6

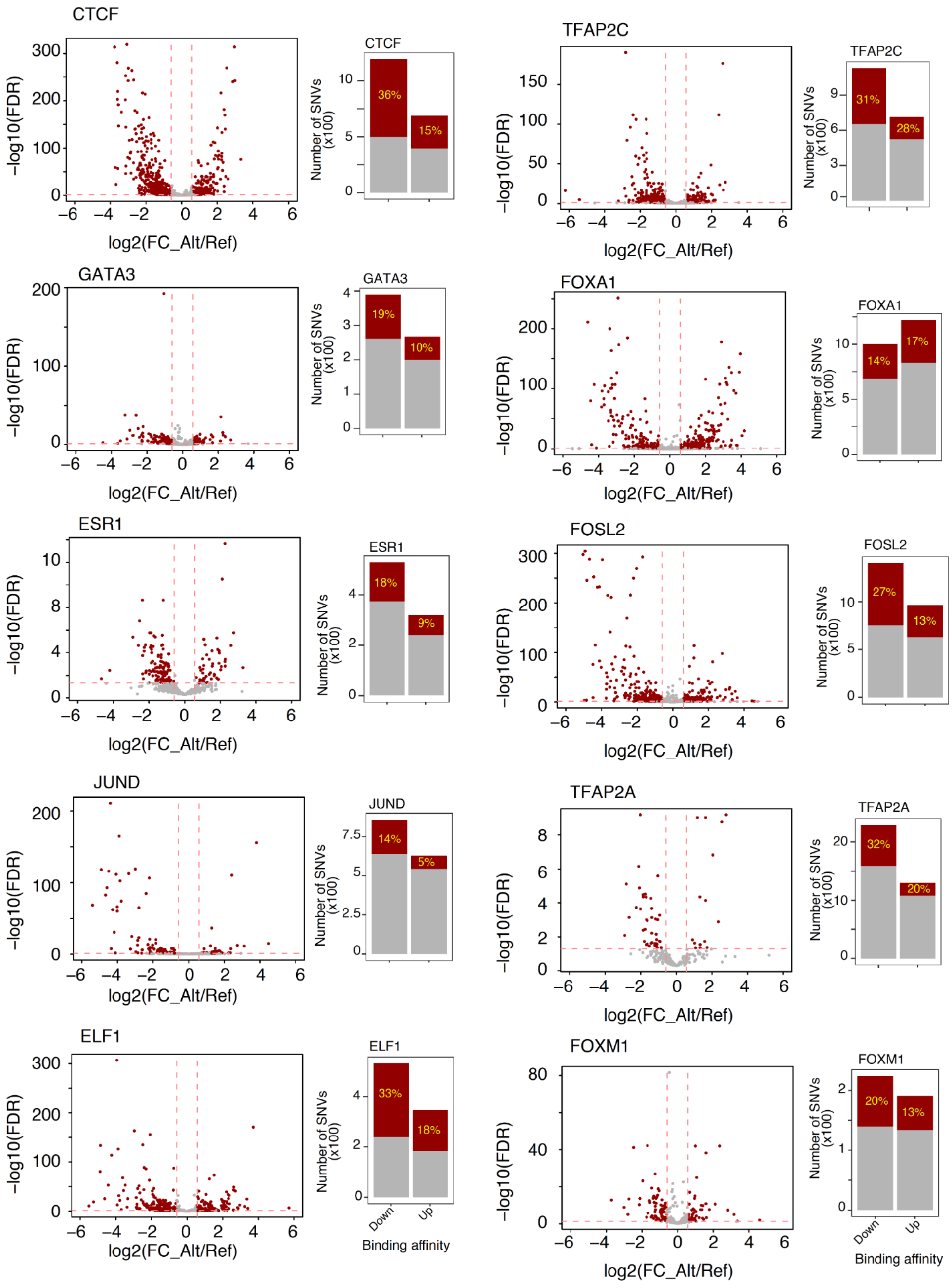


**Figure S6: Mutations within specific cistromes are predicted to affect binding affinity of transcription factors.** The volcano plots show Bonferroni adjusted p-values (log10(FDR)) and the fold change in binding intensity resulting from somatic mutations within binding sites +/- 100bp. Red dotted lines indicate a p-value ≤ 0.05 and fold change ≥ 1.5. The barplots show the number of mutations that are predicted to affect the binding affinity of specific transcription factors (Up: increased binding affinity; down: decreased binding affinity).

### Supplementary Tables

**Table S1: Princess Margaret cohort**. list of tumors in the initial cohort (n=26).

**Table S2: Signal matrix of accessible chromatin in PM_Lum breast tumors.**

**Table S3: Motif enrichment analysis using as a background DNaseI hypersensitive regions and target regions from** (**S3.1**) PM_Lum and (**S3.2**) TCGA_Lum.

**Table S4: Summary of mutation calls from 16 different cancer types overlapping PM_Lum Catalogue**

**Table S5: Results of mutation enrichment analysis using ActiveDriverWGS and our in-house algorithm**.
